## Supplemental material for "Chromatin Environment-Dependent Effects of DOT1L on Gene Expression in Male Germ Cells"

Fig. S1

a

H3K79me2 ChIP-Seq

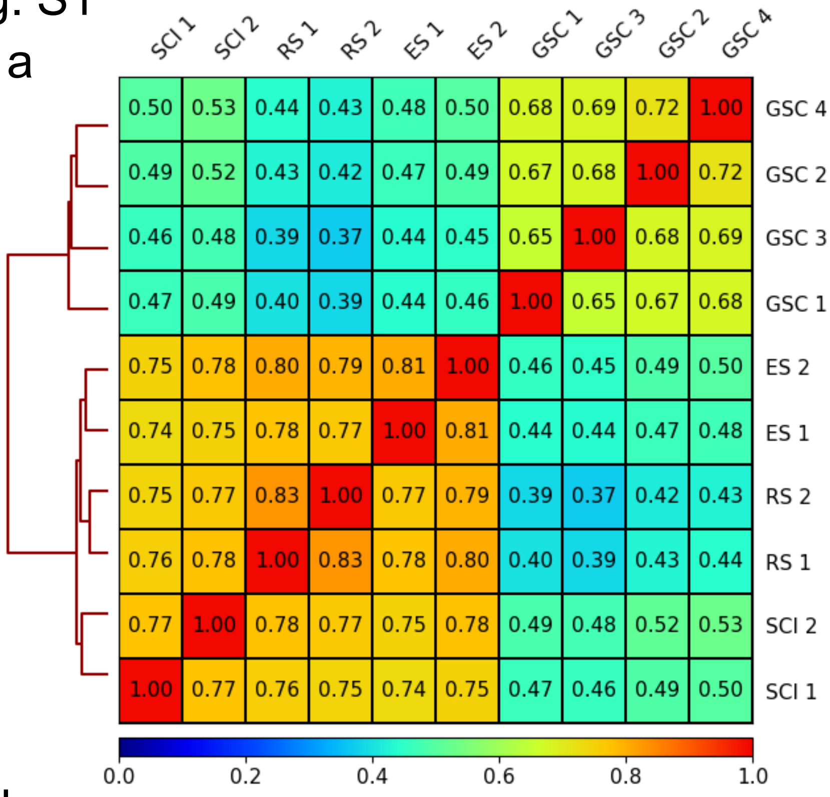

b

| | GSC rep1 | GSC rep2 | GSC rep3 | GSC rep4 | SCI rep1 | SCI rep2 | RS rep1 | RS rep2 | ES rep1 | ES rep2 |
|---|---|---|---|---|---|---|---|---|---|---|
| Number of reads | 21 260 743,00 | 23 051 274,00 | 24 537 676,00 | 25 195 852,00 | 45 210 180,00 | 54 045 728,00 | 56 859 200,00 | 53 051 834,00 | 43 379 619,00 | 55 919 709,00 |
| **MACS2 peak calling (broad parameter)** | | | | | | | | | | |
| Number of peaks | 26 856,00 | 26 463,00 | 25 571,00 | 28 820,00 | 53 603,00 | 53 895,00 | 70 458,00 | 75 418,00 | 7 678,00 | 1 276,00 |
| Number of annotated peaks | 26 601,00 | 26 189,00 | 25 325,00 | 28 505,00 | 52 977,00 | 53 292,00 | 69 485,00 | 74 382,00 | 7 607,00 | 1 225,00 |
| **common analysis** | | | | | | | | | | |
| Number of peaks in common | | | | 15 896,00 | 46 034,00 | | 67 918,00 | | | 385,00 |
| **ChIPseeker** | | | | | | | | | | |
| Number of peaks annotated as "promoter" | | 10 089,00 | | | 19 480,00 | | 25 555,00 | | 4 040,00 | |
| Number of peaks annotated as "intragenic" | | 5 621,00 | | | 13 880,00 | | 21 348,00 | | 4 267,00 | |
| Number of peaks annotated as "distal intergenic" | | | | 170,00 | 2 122,00 | | 8 788,00 | | | 159,00 |
| Number of peaks annotated as "downstream" | | | | 15,00 | | 219,00 | | 535,00 | | 7,00 |

c

H3K27ac CUT&Tag

H3K27me3 CUT&Tag

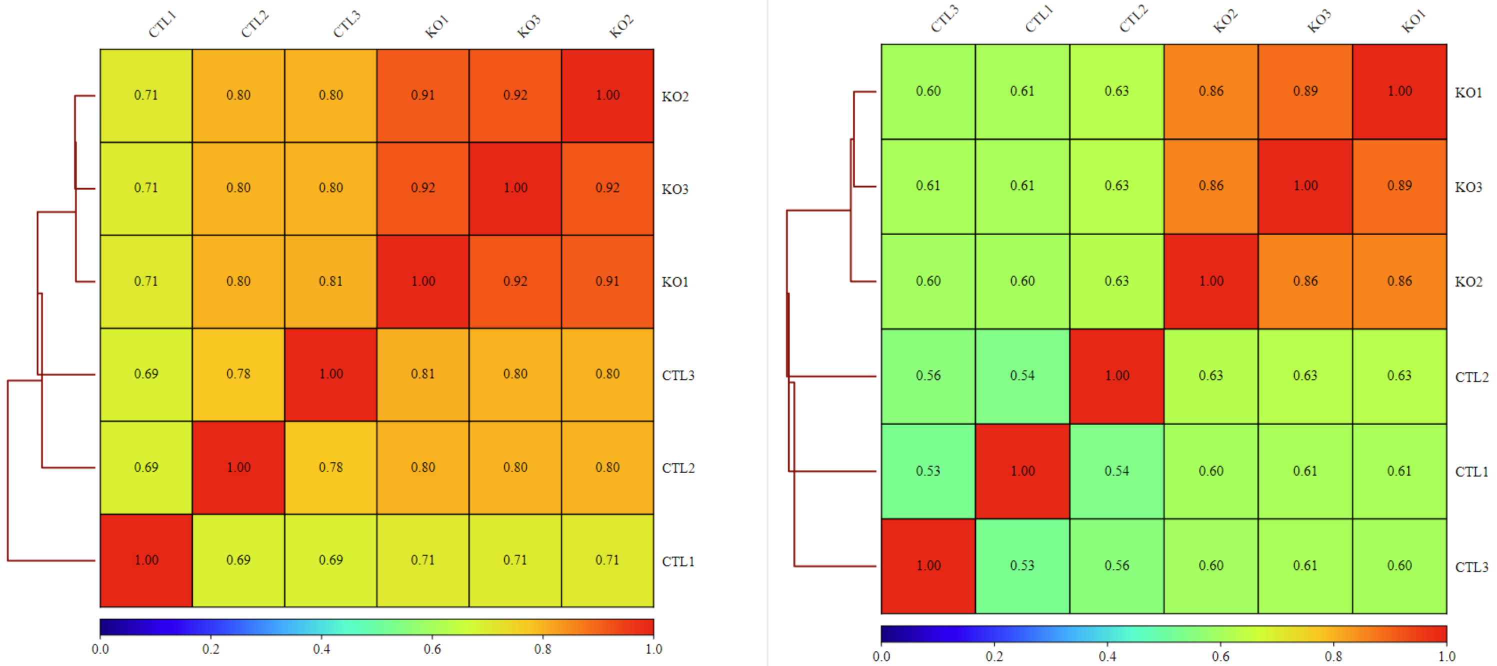

Fig. S2

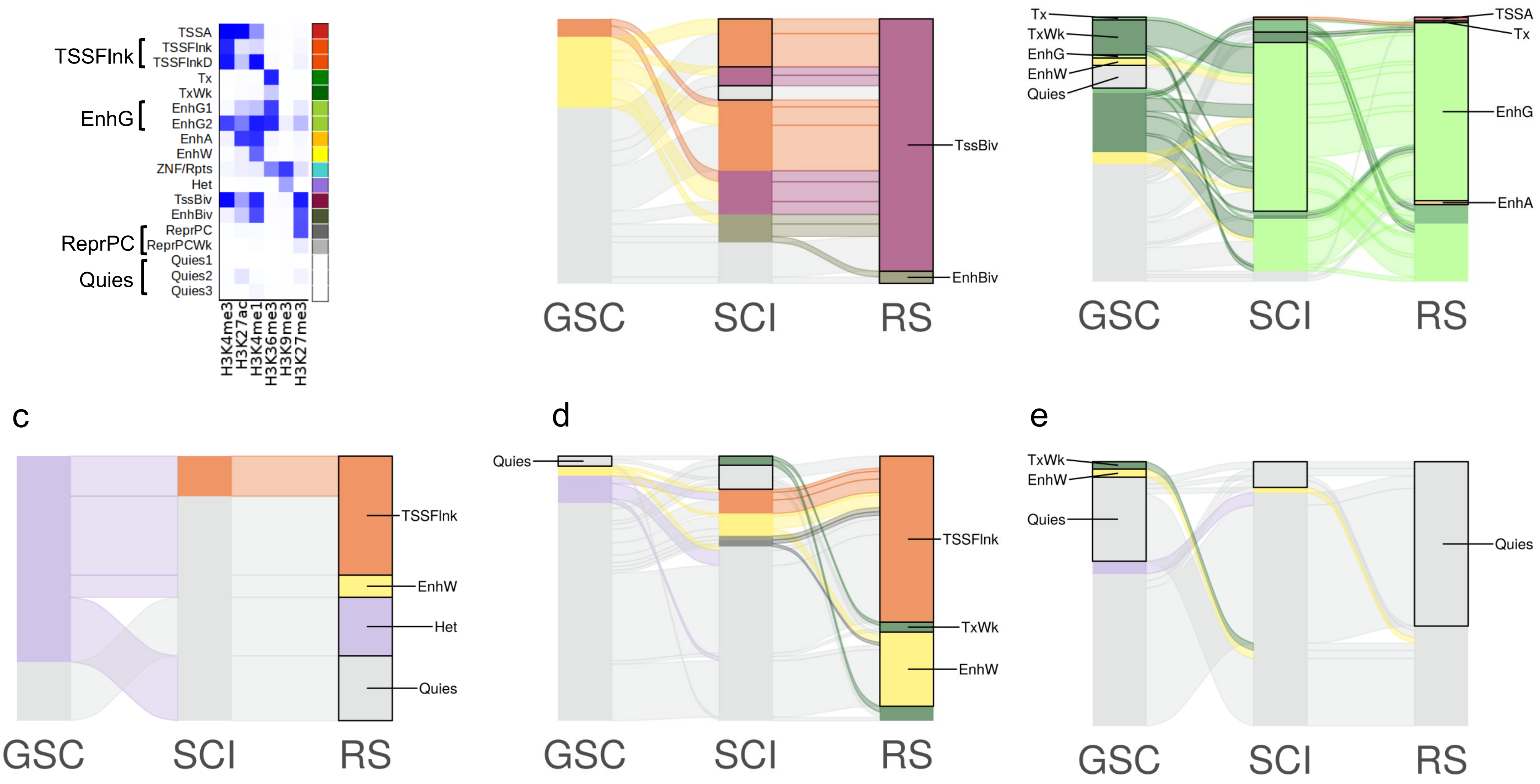
TSSFlnk
EnhG
ReprPC
Quies
TSSA
TSSFlnk
TSSFlnkD
Tx
TxWk
EnhG1
EnhG2
EnhA
EnhW
ZNF/Rpts
Het
TssBiv
EnhBiv
ReprPC
ReprPCWk
Quies1
Quies2
Quies3
H3K4me3
H3K27ac
H3K4me1
H3K36me3
H3K9me3
H3K27me3
a
TssBiv
EnhBiv
GSC
SCI
RS
b
Tx
TxWk
EnhG
EnhW
Quies
TSSA
Tx
EnhG
EnhA
GSC
SCI
RS
c
TSSFlnk
EnhW
Het
Quies
GSC
SCI
RS
d
Quies
TSSFlnk
TxWk
EnhW
GSC
SCI
RS
e
TxWk
EnhW
Quies
Quies
GSC
SCI
RS

Fig.S3

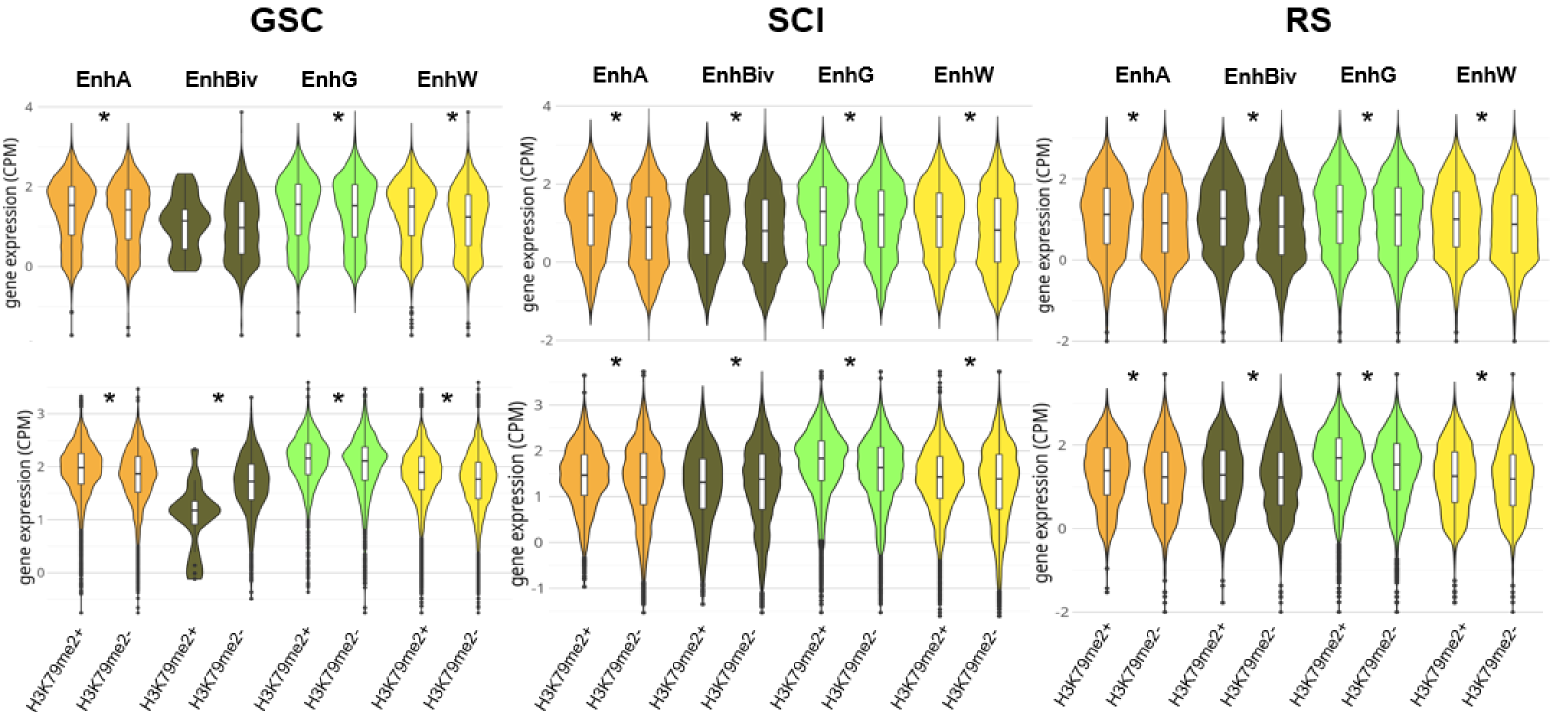
GSC
SCI
RS
EnhA
EnhBiv
EnhG
EnhW
gene expression (CPM)
H3K79me2+
H3K79me2-
*

Fig. S4

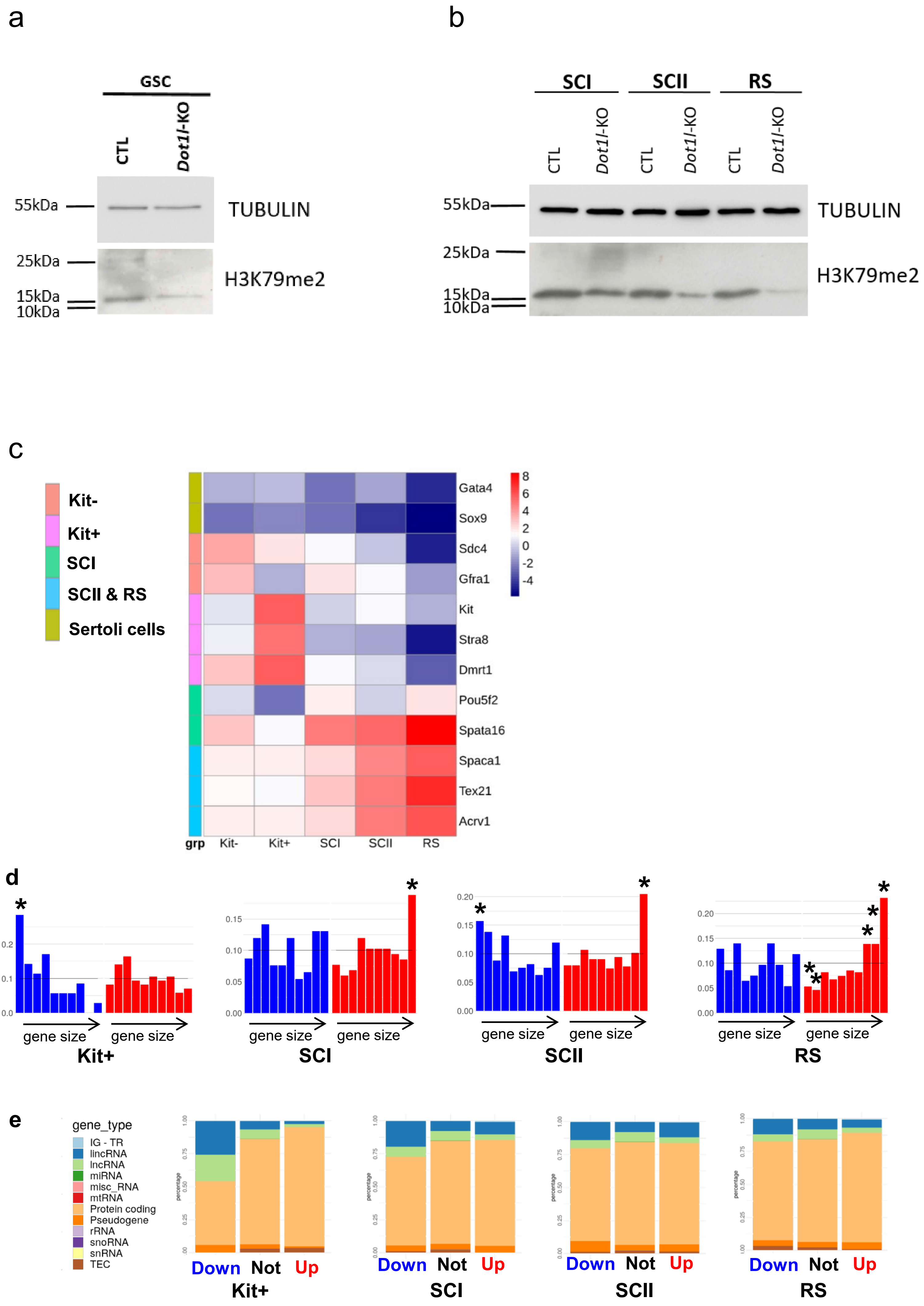

a
GSC
CTL
Dot1l-KO
55kDa
TUBULIN
25kDa
15kDa
10kDa
H3K79me2
b
SCI
SCII
RS
CTL
Dot1l-KO
CTL
Dot1l-KO
CTL
Dot1l-KO
55kDa
TUBULIN
25kDa
15kDa
10kDa
H3K79me2
c
Kit-
Kit+
SCI
SCII & RS
Sertoli cells
Gata4
Sox9
Sdc4
Gfra1
Kit
Stra8
Dmrt1
Pou5f2
Spata16
Spaca1
Tex21
Acrv1
8
6
4
2
0
-2
-4
grp
Kit-
Kit+
SCI
SCII
RS
d
gene size
gene size
Kit+
gene size
gene size
SCI
gene size
gene size
SCII
gene size
gene size
RS
e
gene_type
IG - TR
lincRNA
lncRNA
miRNA
misc_RNA
mtRNA
Protein coding
Pseudogene
rRNA
snoRNA
snRNA
TEC
Down Not Up
Kit+
Down Not Up
SCI
Down Not Up
SCII
Down Not Up
RS

Fig. S5

a

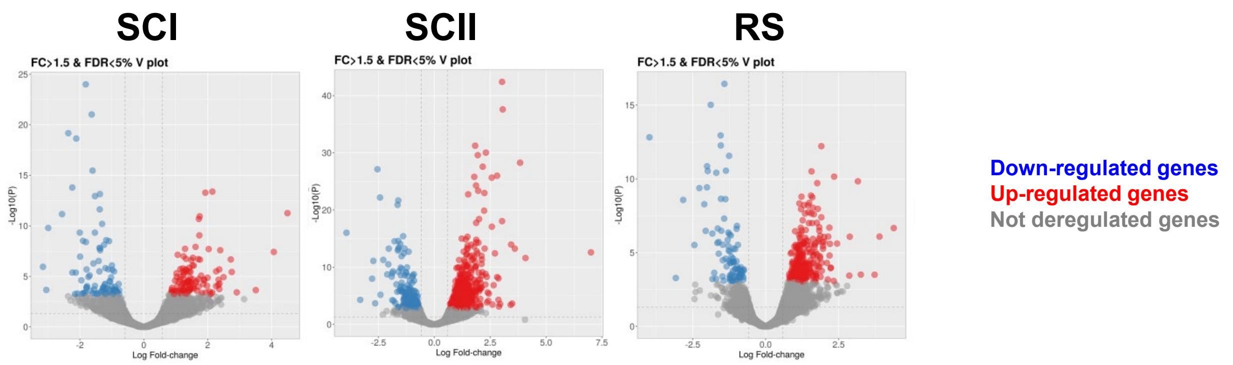
SCI
SCII
RS
FC>1.5 & FDR<5% V plot
-Log10(P)
Log Fold-change
Down-regulated genes
Up-regulated genes
Not deregulated genes

b

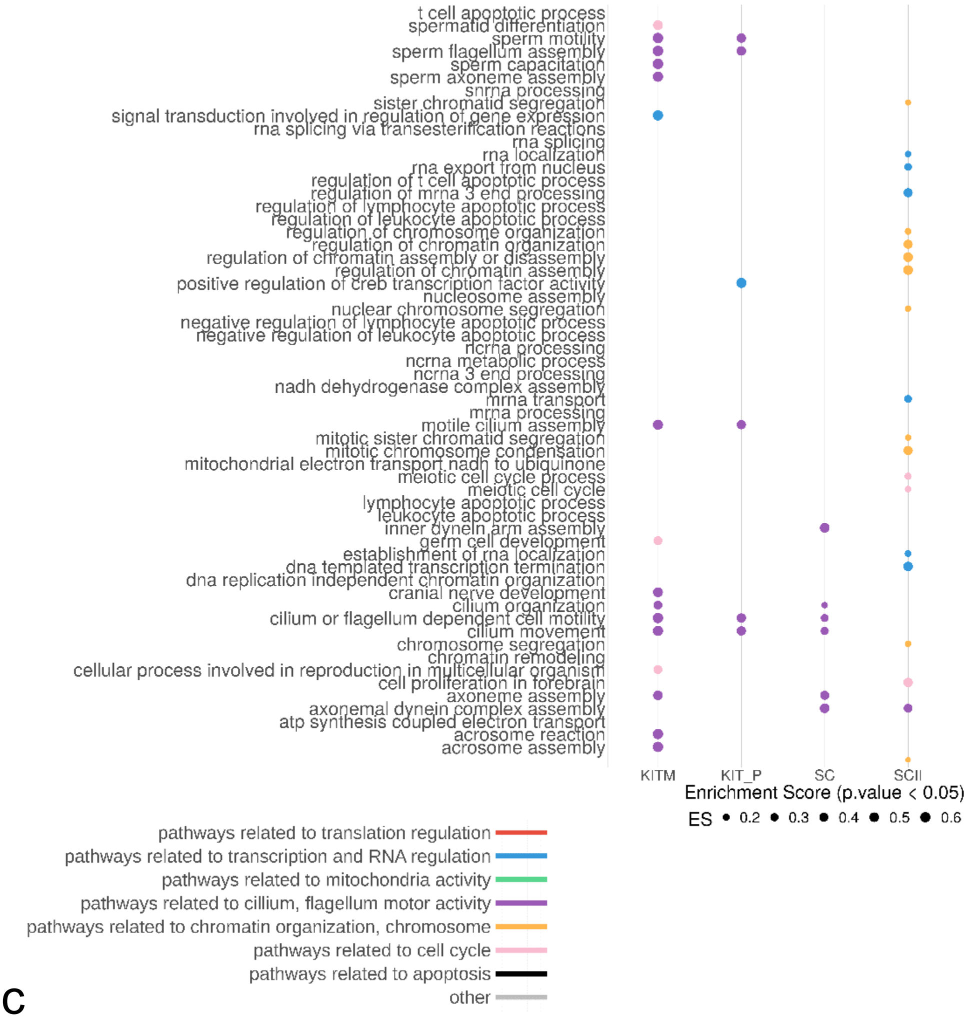
t cell apoptotic process
spermatid differentiation
sperm motility
sperm flagellum assembly
sperm capacitation
sperm axoneme assembly
snrna processing
sister chromatid segregation
signal transduction involved in regulation of gene expression
rna splicing via transesterification reactions
rna splicing
rna localization
rna export from nucleus
regulation of t cell apoptotic process
regulation of mrna 3 end processing
regulation of lymphocyte apoptotic process
regulation of leukocyte apoptotic process
regulation of chromosome organization
regulation of chromatin organization
regulation of chromatin assembly or disassembly
regulation of chromatin assembly
positive regulation of creb transcription factor activity
nucleosome assembly
nuclear chromosome segregation
negative regulation of lymphocyte apoptotic process
negative regulation of leukocyte apoptotic process
ncrna processing
ncrna metabolic process
ncrna 3 end processing
nadh dehydrogenase complex assembly
mrna transport
mrna processing
motile cilium assembly
mitotic sister chromatid segregation
mitotic chromosome condensation
mitochondrial electron transport nadh to ubiquinone
meiotic cell cycle process
meiotic cell cycle
lymphocyte apoptotic process
leukocyte apoptotic process
inner dynein arm assembly
germ cell development
establishment of rna localization
dna templated transcription termination
dna replication independent chromatin organization
cranial nerve development
cilium organization
cilium or flagellum dependent cell motility
cilium movement
chromosome segregation
chromatin remodeling
cellular process involved in reproduction in multicellular organism
cell proliferation in forebrain
axoneme assembly
axonemal dynein complex assembly
atp synthesis coupled electron transport
acrosome reaction
acrosome assembly
KITM
KIT_P
SC
SCII
RS
Enrichment Score (p.value < 0.05)
ES 0.2 0.3 0.4 0.5 0.6
pathways related to translation regulation
pathways related to transcription and RNA regulation
pathways related to mitochondria activity
pathways related to cillium, flagellum motor activity
pathways related to chromatin organization, chromosome
pathways related to cell cycle
pathways related to apoptosis
other

c

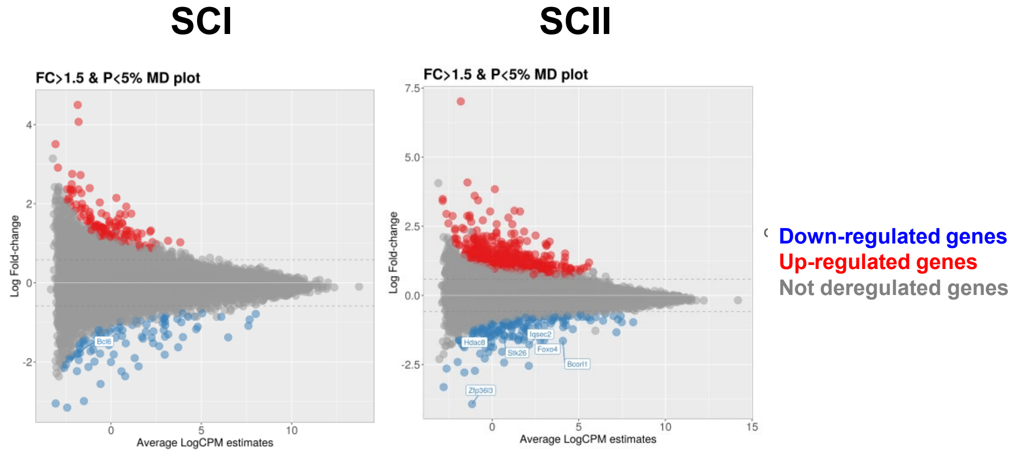
SCI
SCII
FC>1.5 & P<5% MD plot
Log Fold-change
Average LogCPM estimates
Down-regulated genes
Up-regulated genes
Not deregulated genes

Fig. S6

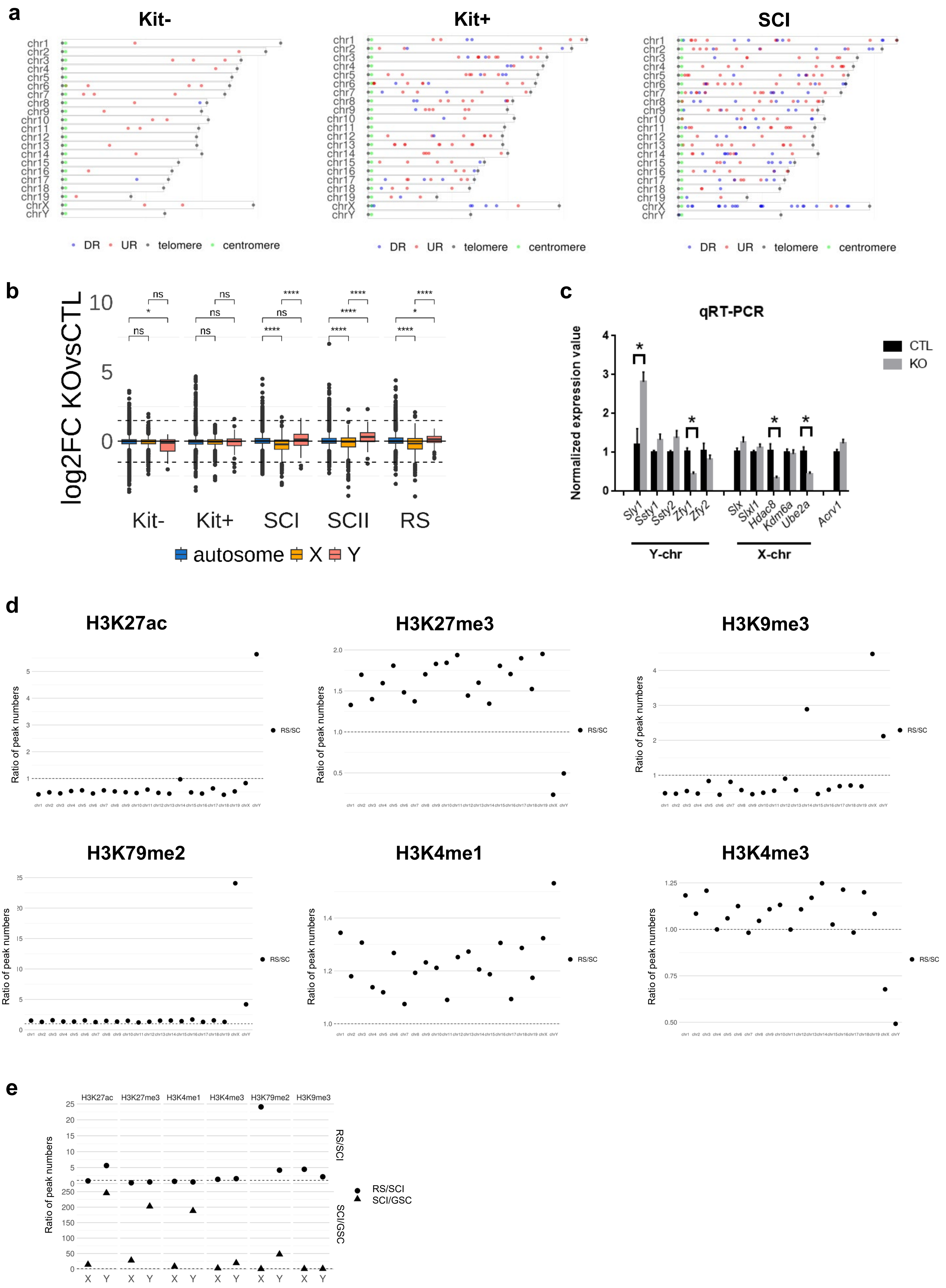
a
Kit-
Kit+
SCI
chr1
chr2
chr3
chr4
chr5
chr6
chr7
chr8
chr9
chr10
chr11
chr12
chr13
chr14
chr15
chr16
chr17
chr18
chr19
chrX
chrY
DR
UR
telomere
centromere
b
log2FC KOvsCTL
10
5
0
ns
*
****
Kit-
Kit+
SCI
SCII
RS
autosome
X
Y
c
qRT-PCR
Normalized expression value
CTL
KO
Sly1
Ssty1
Ssty2
Zfy1
Zfy2
Slx
Slxl1
Hdac8
Kdm6a
Ube2a
Acrv1
Y-chr
X-chr
d
H3K27ac
H3K27me3
H3K9me3
H3K79me2
H3K4me1
H3K4me3
Ratio of peak numbers
RS/SC
e
H3K27ac
H3K27me3
H3K4me1
H3K4me3
H3K79me2
H3K9me3
RS/SCI
SCI/GSC
X
Y

Fig. S7

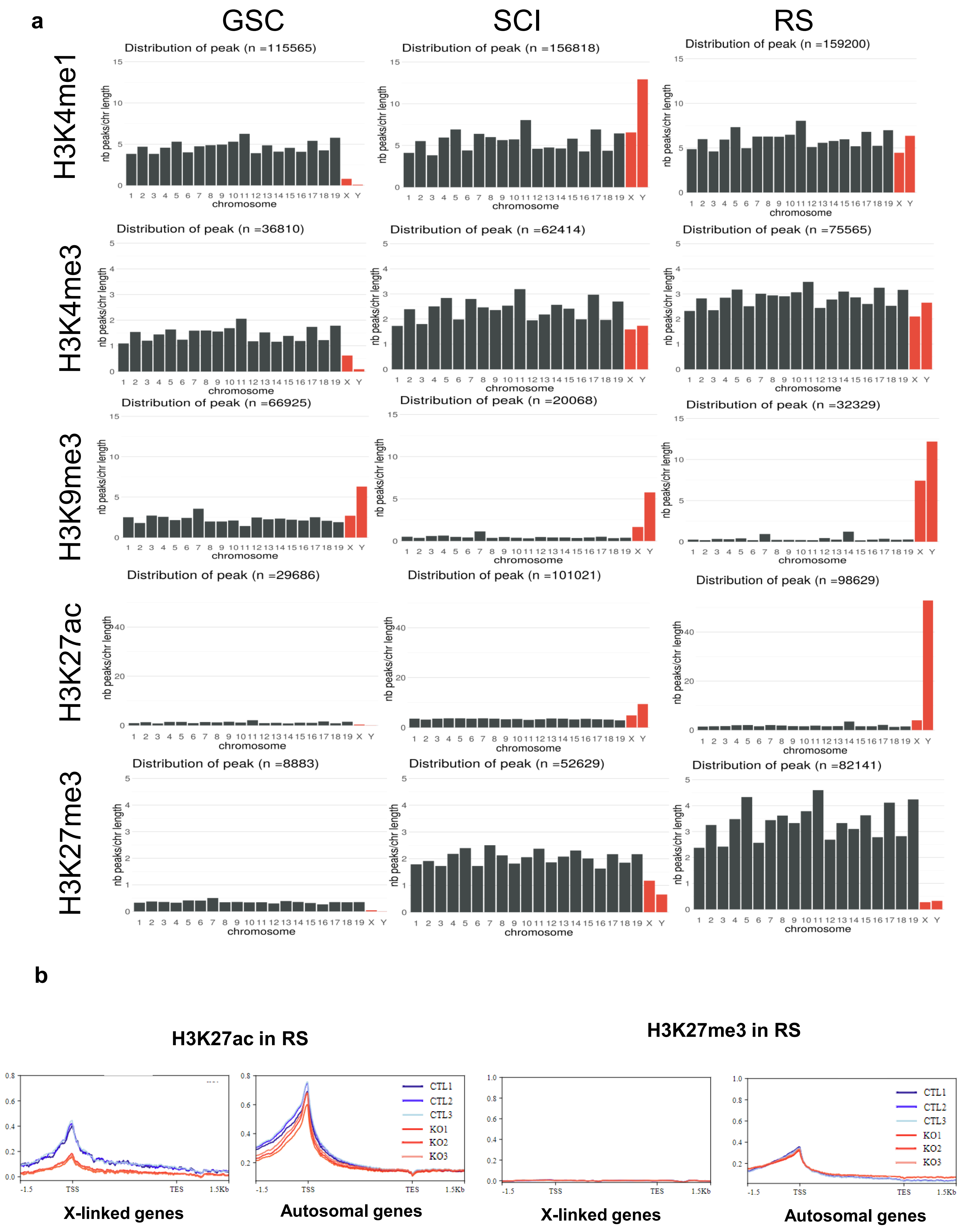
a
GSC
SCI
RS
H3K4me1
H3K4me3
H3K9me3
H3K27ac
H3K27me3
Distribution of peak (n =115565)
Distribution of peak (n =156818)
Distribution of peak (n =159200)
Distribution of peak (n =36810)
Distribution of peak (n =62414)
Distribution of peak (n =75565)
Distribution of peak (n =66925)
Distribution of peak (n =20068)
Distribution of peak (n =32329)
Distribution of peak (n =29686)
Distribution of peak (n =101021)
Distribution of peak (n =98629)
Distribution of peak (n =8883)
Distribution of peak (n =52629)
Distribution of peak (n =82141)
nb peaks/chr length
chromosome
b
H3K27ac in RS
H3K27me3 in RS
CTL1
CTL2
CTL3
KO1
KO2
KO3
TSS
TES
X-linked genes
Autosomal genes

Fig. S8

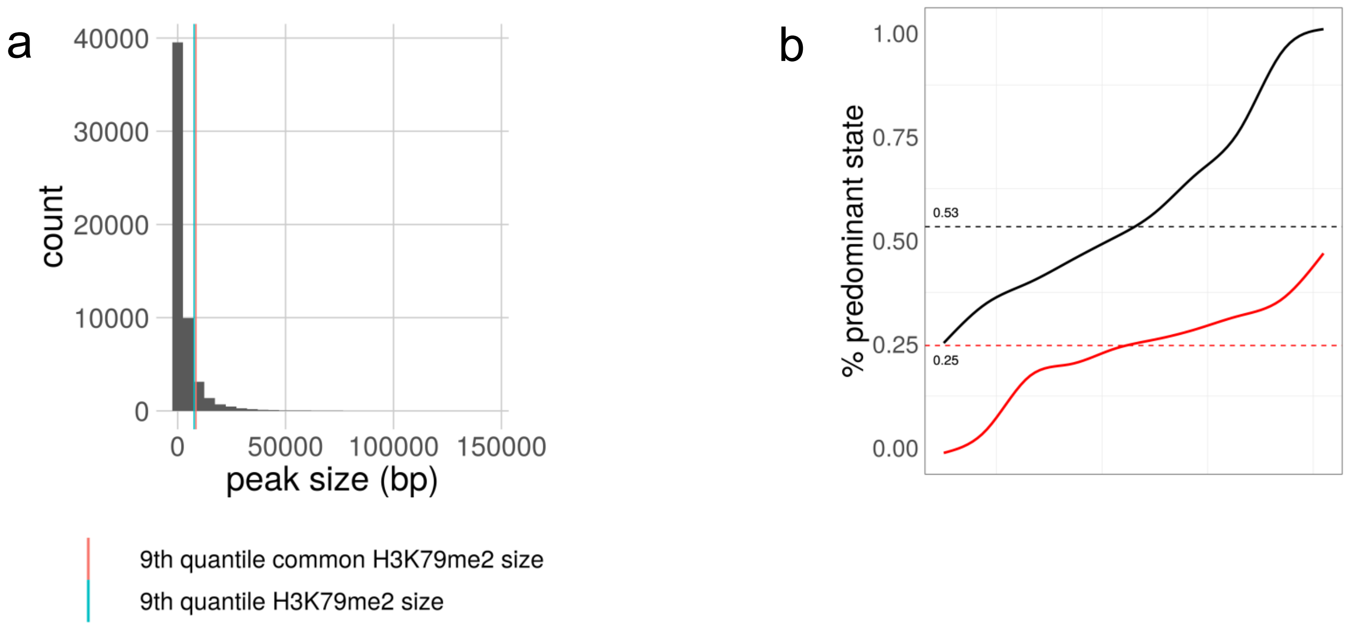
a
40000
30000
20000
10000
0
count
0
50000
100000
150000
peak size (bp)
9th quantile common H3K79me2 size
9th quantile H3K79me2 size
b
1.00
0.75
0.50
0.25
0.00
% predominant state
0.53
0.25
